# Arthropod community composition in urban landscapes is shaped by both environmental filtering and dispersal limitation

**DOI:** 10.1101/2024.01.09.574897

**Authors:** Indigo R. Roper-Edwards, Allen H. Hurlbert

## Abstract

We assessed the relative importance of environmental filtering and dispersal limitation in structuring foliage- and ground-dwelling arthropod communities in central North Carolina. We hypothesized that both the local environment and the dispersal distance between sites would predict community composition, but that dispersal distance would be more important for ground arthropods than for foliage arthropods. In both groups, variation in habitat characteristics was important in structuring communities, and the role of dispersal in structuring communities was much greater after accounting for variation in landscape connectivity. Our results demonstrate the importance of both dispersal limitation and environmental filtering in shaping community composition and emphasize the importance of variation in the landscape for modeling these forces. Examining communities of multiple arthropod groups across the same spatial gradient highlights the scale-dependence of these processes and illustrates how variation in the environment can alter the relative abundance of specialist and generalist taxa.

## Introduction

Landscape change due to human activity poses threats to ecosystems worldwide. Urbanization, in particular, modifies ecosystems at both the landscape scale, through habitat loss and fragmentation, and at the local scale, through changes in the environmental conditions. These changes can include increases in temperature, artificial light, and impervious surface area, as well as changes to plant communities and the density of edges between habitat and non-habitat [1–6]. Although the varied components of urbanization present a challenge for identifying the mechanisms by which it impacts ecosystems, doing so is essential for protecting those systems as human populations continue to grow.

In broad terms, the four high-level processes that shape community composition as classified by Vellend [7,8] are natural selection, dispersal, speciation, and ecological drift. While speciation may occasionally be associated with urbanization [9,10], it plays a relatively minor role over the spatial and temporal scales of urban community assembly. Ecological drift, or the random variation in the abundance of species unattributable to species traits, can lead to differences in community composition between sites but is, by definition, unpredictable, and it manifests as unexplained variation. This leaves selection and dispersal as the two vital processes by which environmental changes associated with urbanization alter the composition of communities across the landscape, and as such, we chose to investigate the relative importance of these two forces. Specifically, we set out to characterize the roles of environmental filtering, or selection based on the local environment, and dispersal limitation, often resulting from habitat loss or fragmentation, on community composition.

Selection results from fitness differences between individuals which can arise due to biotic and abiotic forces. Interspecific interactions may vary across an urbanization gradient, but change in environmental conditions is the selective factor most clearly associated with urbanization [2,3]. As such, we focus on the impacts of environmental filtering on community composition. When environmental filtering acts strongly at a site, the composition of that site is assumed to be a subset of the regional species pool, “filtered” to those species which are best suited to the local environment. Therefore, communities in distinct local environments are expected to have distinct compositions [7,11]. For environmental filtering to be solely responsible for shaping a local community, however, all of the species in the regional pool must be capable of colonizing the site.

If species are not able to disperse between one community and another, this limitation, rather than environmental traits, may explain the presence or absence of those species from certain communities. At large scales where many species are involved, the geographic distance, or the effective distance accounting for connectivity, between communities can be a stronger determinant of differences in their composition than the environment in which they occur [12]. We can gain insight into the relative importance of these forces in landscapes altered by urbanization by comparing the relative importance of variables associated with dispersal limitation and environmental filtering in structuring communities [13].

Arthropods are a valuable taxonomic group for assessing the effects of anthropogenic change. They are found across the planet, they are essential components of many ecosystems, and they are responsive to environmental changes. Urbanization has been shown to affect arthropods in particular, manifesting as changes to biodiversity, abundance, and community composition [3,5,14–16]. Arthropod communities are expected to differ across the landscape due to both environmental filtering and dispersal limitation, which modify the species present at any particular location [13,17]. Therefore, we can use arthropods as a model in order to identify the aspects of urbanization that modify community composition, either via environmental filtering or dispersal limitation.

To assess the relative importance of environmental filtering and dispersal limitation in shaping arthropod communities, we characterized the composition of foliage- and ground-dwelling arthropod communities at 30 plots across six sites in central North Carolina. These plots varied in environmental conditions and in their distance from one another. This allowed for a comparison between the effects of dispersal limitation and environmental filtering across two different groups within the same landscape which differ in their habitat requirements and dispersal abilities. We hypothesized that 1) differences in both the local environment and the distance or connectivity between communities would predict compositional differences, 2) ground arthropods would have lower average dispersal ability and therefore their community composition would be more strongly predicted by dispersal limitation, and 3) differences in arthropod community composition between plots would be better predicted by distance measures which account for variation in landscape connectivity than by straight-line distances.

## Materials and Methods

### Sample Collection and Identification

Arthropods were collected from six sites in Orange and Wake Counties in North Carolina (Fig. 1). Each site contained five sampling plots, and sites were selected to span a range of percent forest cover (Table 1) and to capitalize on existing arthropod monitoring sites established for the *Caterpillars Count!* citizen science project [18]. We characterized plots by forest cover, as it represents habitat relevant for foliage and ground arthropods, and in the study area, it is strongly negatively correlated with impervious surface. The five plots at each site were at least 100 meters apart. Arthropod communities were characterized at the scale of each plot.

**Fig. 1:**
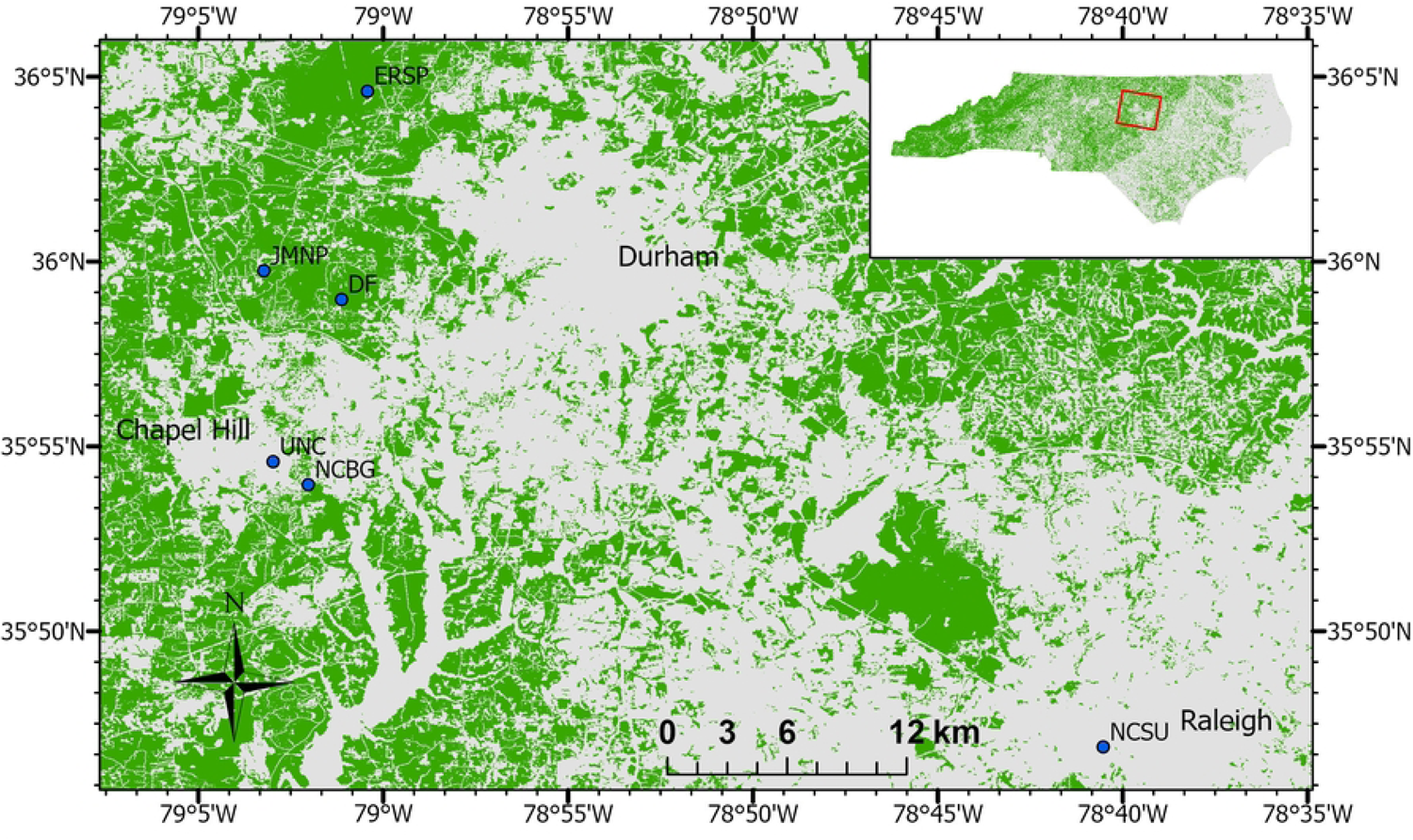
Map of arthropod sampling sites in central North Carolina (blue points). Forested areas as classified in the 2019 National Land Cover Dataset shown in green. Site abbreviations as in Table 1.

**Table 1:** Environmental characteristics (site mean, with range of values across the five plots in brackets) of the six sampling sites.

| Site Name | Site Code | Proportion Forest Cover in a 1-km Radius | Litter Layer Depth (mm) | Percent Canopy Cover | Herbaceous Cover Class (1-4) |
| --- | --- | --- | --- | --- | --- |
| Eno River State Park | ERSP | 0.927 [0.909-0.947] | 26 [15-40] | 77.2 [69-83] | 2.2 [1-3] |
| Duke Forest, Korstian Division | DF | 0.865 [0.844-0.899] | 24 [10-40] | 82.2 [79-86] | 1.6 [1-2] |
| Johnston Mill Nature Preserve | JMNP | 0.816 [0.798-0.836] | 28 [20-40] | 79.8 [77-89] | 1.6 [1-2] |
| North Carolina Botanical Garden | NCBG | 0.420 [0.391-0.445] | 27 [15-40] | 77.6 [71-83] | 1.8 [1-3] |
| North Carolina State University | NCSU | 0.005 [0.001-0.011] | 20 [5-50] | 84.2 [81-87] | 1.4 [1-3] |
| University of North Carolina | UNC | 0.082 [0.072-0.098] | 32 [10-50] | 74 [43-93] | 2.4 [1-4] |

To survey ground arthropods, a pitfall trap was deployed at each of the five plots at each site, following the design recommended by Montgomery et al. [19]. Each trap consisted of a cup, buried flush with the ground, filled with ∼75 ml propylene glycol and with a roof suspended on pegs 4-8 centimeters above the lip of the cup. Each of these traps was active for a total of three 4-day periods from May to July of 2022. The contents of the trap were collected and the propylene glycol replaced after the first two days of each collection period. Arthropods collected in pitfall traps were returned to the lab, strained from the propylene glycol, and preserved in 70% ethanol in plastic vials labeled with the trap and date of collection.

To survey foliage arthropods, beat sheet surveys were performed on a set of five trees at each of five plots per site. Trees were chosen based on the presence of branches with at least fifty leaves at a height of less than two meters from the ground, with tree species being chosen that were representative of the composition of the understory at each site. Beat sheet surveys were performed using the method described for *Caterpillars Count!* sampling by Hurlbert al. [18], with the additional step of transferring arthropods to vials of 70% ethanol labeled with the date and a unique tree identifier. Foliage surveys were conducted during three collection periods at each site between May and July of 2022. Each collection period included three sets of beat sheet surveys, two days apart.

To measure biomass, foliage and ground arthropods were removed from ethanol and allowed to dry for about 5 minutes, then weighed. Once arthropods were returned to the lab, they were identified to the family level, with the exception of ants (family Formicidae), which were identified to genus [20]. Arthropods were identified using books and online resources [21–25]. After identification, we retrieved the unique ID number and taxonomic information for each observed taxon from the Integrated Taxonomic Information System (ITIS) in order to ensure standardized taxonomic nomenclature. For taxa not recognized by ITIS, up-to-date taxonomic information was verified using BugGuide [26], or, in the case of the ant species *Brachyponera chinensis*, AntWiki. For these taxa, unique taxon IDs were individually assigned.

Several environmental variables were measured at each plot. Litter depth was measured in millimeters using a marked skewer, local canopy cover was estimated from photos taken using a cell phone camera directed vertically from ground level, and a categorical estimate of herbaceous plant cover within a 1-meter radius of the pitfall trap was recorded as 0-25%, 26-50%, 51-75%, or 76-100% cover. Finally, for each sampling plot, the proportion of forest cover within a 1 kilometer radius was calculated from a 30-meter resolution National Land Cover Database [27] raster image from 2019. It should be noted that the proportion forest cover values from plots at the same site are not entirely independent, as plots were generally less than 1 kilometer apart.

Two different measurements of distance between sampling plots were calculated, geographic distance and resistance of the shortest path (hereafter, resistance), where resistance represented the effective dispersal distance between a pair of sampling plots where the landscape between those plots varied in permeability for dispersal [28]. In theory, resistance effectively accounts for fragmented landscapes, where the geographic distance between two points might be short, but an intervening area of inhospitable habitat impedes direct dispersal. To generate a resistance landscape, we reclassified the NLCD land cover raster to resistance values from 1 (greatest permeability for dispersal) to 10 (greatest resistance to dispersal). Because this study focused on forest arthropods, all forest classes were assigned a value of 1, all developed cover classes except developed open space were assigned a value of 10, all water and barren areas were assigned a value of 10, and all non-forest plant cover, including agriculture and developed open space, were assigned a value of 5. We explored alternative resistance landscapes that ranked cover classes in similar order but that used different numerical resistance values for each class, but found strong correlations among the resulting resistance of shortest path values, likely because urban and forest land cover categories accounted for the majority of the landscape (*R^2^* > 0.94; Fig. S1). The reclassified raster was used with CircuitScape [29], along with the coordinates of the sampling plots, converted into the coordinate system of the raster, to generate the total resistance of the least-resistance path between each pair of sampling plots.

### Data Analysis

We characterized the dissimilarity of arthropod communities between sample plots using two complementary metrics. Euclidean distance between sampling plots *x* and *y* (*E_xy_*) was calculated using the biomass of arthropods in each family observed at a plot as:

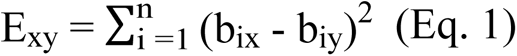

where *b_ix_* and *b_iy_* are the log-transformed values of biomass of family *i* at sampling plots *x* and *y*, respectively. Biomass values were log-transformed to prevent the most abundant or large-bodied arthropod families from having an outsized impact on Euclidean distance, as the biomass of different arthropod families can vary by several orders of magnitude. Euclidean distance between plots is still primarily influenced by the differences in biomass of common families.

To capture differences in community composition driven by the gain or loss of species irrespective of differences in biomass, we calculated the Jaccard dissimilarity index (*J_xy_*), which is simply the proportion of families unique to either plot (*f_unique_*) out of the total number of families observed at either plot (*f_total_*):

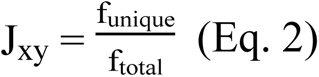

Because this is a presence/absence-based metric, it is unrelated to differences in the biomass of each family between two sampling plots. Therefore, Jaccard distance more effectively captures differences in the presence of rare families between plots. Each metric was calculated separately for ground and foliage arthropod communities.

We evaluated the extent to which dissimilarity in community composition between two plots could be explained by the difference in environmental conditions, by geographic distance, or by resistance of the least-resistance path between those two plots. For foliage arthropod communities, we examined differences in local canopy cover, proportion of forest cover within a 1-km radius, and the Jaccard dissimilarity of sampled tree species between plots. For ground arthropod communities, we examined differences in proportion forest cover, herbaceous plant cover class, and litter depth. Because of the non-independence of all pairwise comparisons, we used Mantel tests to assess the significance of correlations between distance matrices. We used linear models to visualize relationships and to determine the amount of variance in community composition explained by each predictor. Because the site at NC State University differed in proportion of forest cover and distance from other sites to a much greater extent than the other sites, the same analyses were run without the data from this site to determine whether the site had an outsized effect on the observed patterns.

To aid in the interpretation of which families and functional groups contributed most to differences in community composition, principal component analyses were conducted for both foliage and ground arthropods based on log-transformed biomass values at each sampling plot. We classified each arthropod family as predominantly herbivore, omnivore, predator or scavenger with regard to diet. For families which do not exhibit a consistent tendency in diet, the group was assigned to “mixed.” Each family was also classified by the flight ability of its species, where either none, all, or some of the species in the family are capable of flight.

## Results

Several common trends emerged for both foliage and ground arthropod communities regarding predictors of differences in community composition. For both groups, environmental distance was a stronger predictor of differences in community composition than geographic distance, but resistance of the shortest path explained more variation than geographic distance (Figs. 2, 3). For foliage arthropods, resistance explained more variance than any single environmental variable. Differences in the proportion of forest cover within a 1-km radius were predictive of differences in community composition for both groups, but particularly for ground arthropods, explaining nearly fifty percent of the variance in Euclidean distance of ground arthropod community composition (Figure 3a). While forest cover was important for foliage arthropod communities as well, the Jaccard dissimilarity of sampled tree species was more strongly predictive of differences in these communities, accounting for twelve percent of the variance in Euclidean distance (Fig 2e). Geographic distance explained similarly small amounts of variance in community composition differences for both arthropod groups (R^2^_foliage_= 7-9%, R^2^_ground_= 4-9%; Figure 2g-2h, Figure 3g-3h). However, for foliage arthropods, resistance of the shortest path between plots explained more variation in differences in community composition than any other predictor (R^2^ = 25-26%; Figure 2i-2j). For ground arthropods, resistance explained roughly the same amount of variation in Euclidean distance as geographic distance, but resistance explained three times as much variation in Jaccard dissimilarity as geographic distance. There were no variables that were predictive of differences in community composition measured as Euclidean distance but not as Jaccard distance, or vice versa. When the analyses were run excluding data from NC State University, R^2^ values either remained the same or increased for all variables with the exception of geographic distance for foliage arthropods. Otherwise, all comparisons regarding the relative importance of variables were qualitatively similar.

**Fig. 2:**
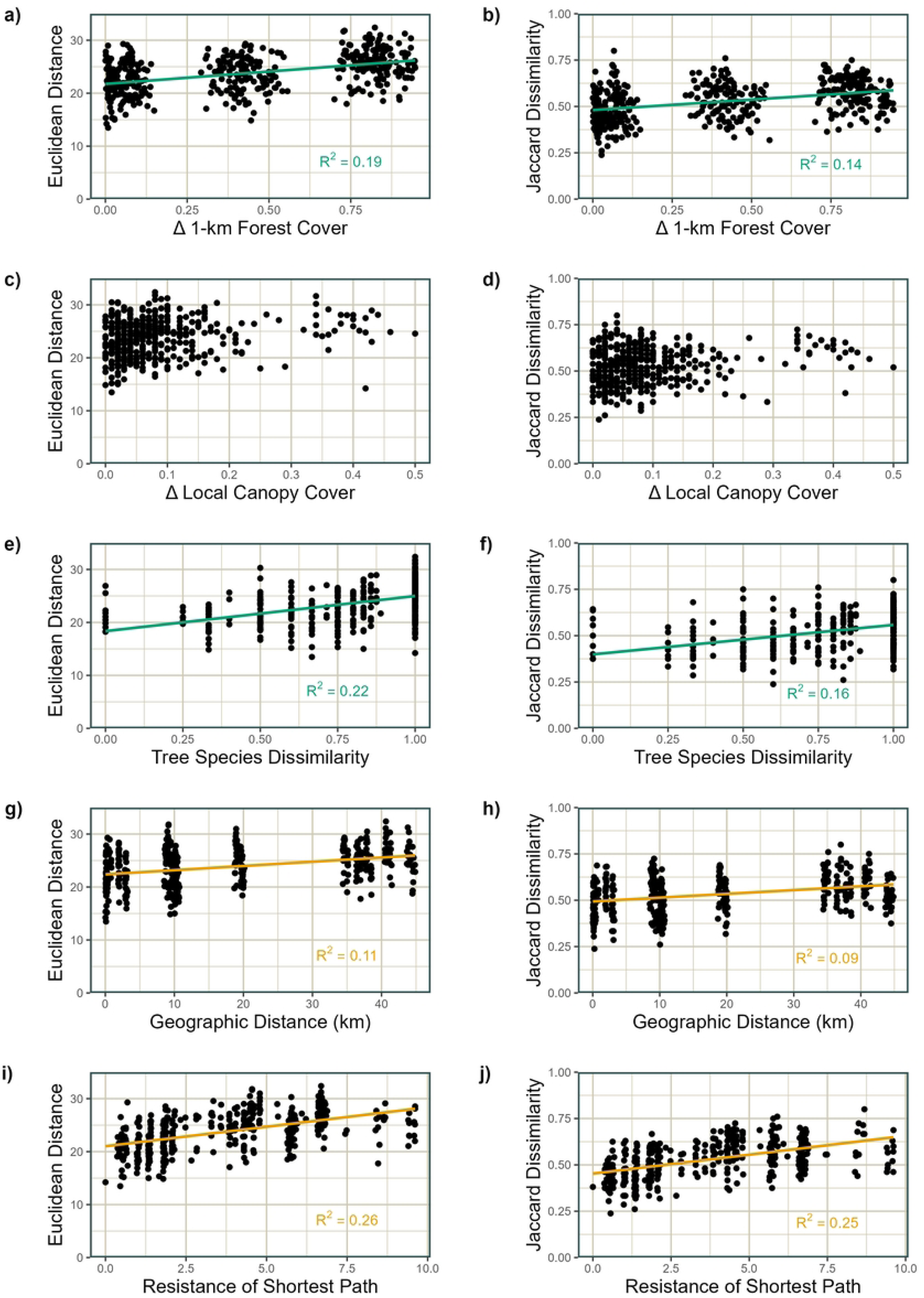
Linear models relating foliage arthropod community composition to environmental predictors. Euclidean distance (a, c, e, g, i) or Jaccard dissimilarity (b, d, f, h, j) of foliage arthropod communities versus (a, b) differences in the proportion of forest cover within 1 km, (c, d) differences in local canopy cover, (e, f) Jaccard dissimilarity of sample tree species, (g, h) geographic distance between sampling plots, and (i, j) resistance of the least-resistance path between sampling plots. Model fits and R^2^ values are displayed only where Mantel test p-values are less than 0.005.

**Fig 3:**
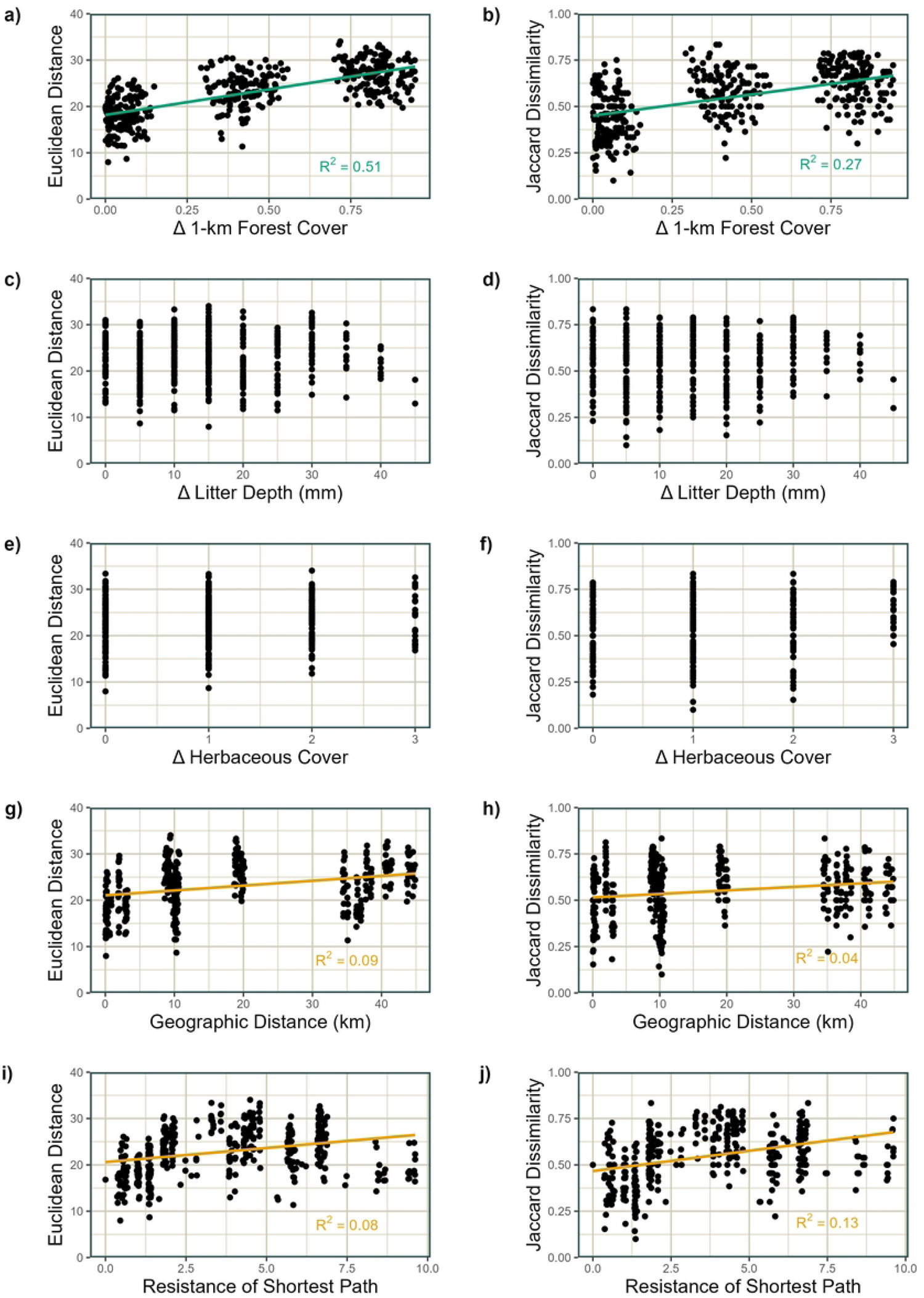
Linear models relating ground arthropod community composition to environmental predictors. Euclidean distance (a, c, e, g, i) or Jaccard dissimilarity (b, d, f, h, j) of ground arthropod communities versus (a, b) differences in the proportion of forest cover within 1 km, (c, d) differences in local litter layer depth, (e, f) differences in herbaceous cover class, and (g, h) geographic distance between sampling plots, and (i, j) resistance of the least-resistance path between sampling plots. Model fits and R^2^ values are displayed only where Mantel test p-values are less than 0.005.

In the principal component analyses for both foliage and ground arthropods, PC1 was strongly correlated with the proportion of forest cover in a 1-km radius (R^2^ > 0.7), while PC2 was not correlated with any of the environmental variables included in our study for either foliage or ground arthropods. For foliage arthropods, lady beetles (Coccinellidae) are associated with lower forest cover, while orb weavers (Araneidae) and sclerosomatid harvestmen, both non-flying families, are associated with greater forest cover. Darkling beetles (Tenebrionidae), also non-flying, have a strong positive loading on PC2. For ground arthropods, only porcellionid woodlice were strongly associated with lower forest cover (Fig. 4a). Camel crickets (Rhaphidophoridae), ground beetles (Carabidae), and wolf spiders (Lycosidae) were all associated with greater forest cover, but ground beetles had a very large positive loading on PC2, while wolf spiders had a relatively strong negative loading on PC2 (Fig. 4b). All of these families are non-flying.

**Fig. 4:**
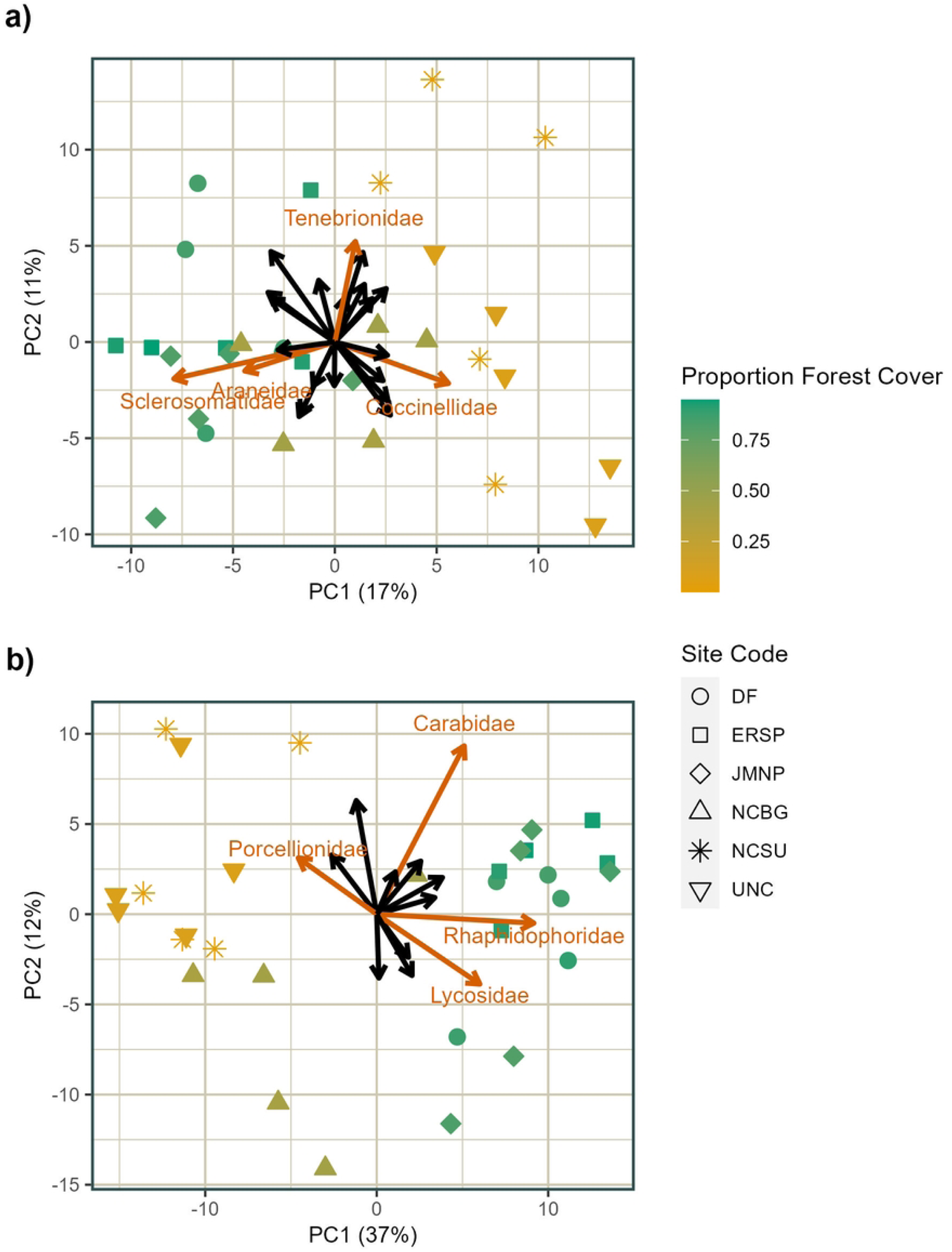
Principal component analysis of foliage and ground arthropod communities. Foliage arthropods (a) and ground arthropods (b), with loadings displayed as arrows and sampling plot values marked with symbols according to site, color-coded by the proportion of forest cover in a 1-km radius. Loadings for families mentioned in the text are labeled and highlighted in color. Site codes correspond to those from Table 1.

## Discussion

Arthropod community composition varied predictably for both foliage and ground arthropods. As hypothesized, environmental differences between plots more strongly predicted compositional differences than the geographic distance between them, and resistance of the shortest path between plots generally predicted differences in the communities of those plots more strongly than the geographic distance between them. In fact, for foliage arthropods, resistance of the shortest path was a stronger predictor than any environmental variable. In contrast to our hypotheses, dispersal limitation, as measured by resistance, had a greater role in determining community composition for foliage arthropods than for ground arthropods. Both the joint role of dispersal limitation and environmental filtering and the greater importance of dispersal in shaping arthropod communities across an urbanization gradient aligns with the findings of Magura et al. [30], who demonstrated that forest arthropod communities are primarily shaped by environmental conditions, while urban arthropod communities are more randomly assembled, suggesting a major role for dispersal limitation.

Traditionally, studies contrasting the compositional effects of environmental filtering with dispersal limitation have used the geographic distance between communities as a proxy for the ease of dispersal between them [12,31]. Our results suggest that this is, at least in some cases, inadequate, as the resistance between sites predicted patterns of community composition change much more strongly than geographic distance. When measured using resistance, dispersal limitation appeared to play a more critical role in shaping foliage arthropod communities, while environmental filtering was the primary force shaping ground arthropod communities. There are two potential interpretations of this finding. One possibility is that, despite the generally greater dispersal ability of foliage arthropods (many of which can fly) compared to ground arthropods, they are less likely to disperse across non-forested areas than ground arthropods. While it is not necessarily true that all foliage arthropods universally have greater dispersal ability than all ground arthropods, this pattern was generally true based on the known flight abilities of the families sampled in our study (Fig. S1). Another possibility is that limitation of ground arthropod dispersal occurs strongly over very short distances, resulting in limited variation to be explained at the landscape level [32]. For example, dissimilarity only appears to increase with distances up to 10 km, after which it is fairly constant (Fig. 3g). This possibility is supported by the results of the analyses when NC State University was excluded - when all of the distances in the analysis were shorter, both geographic distance and resistance more strongly predicted ground arthropod community dissimilarity. To identify the drivers of the patterns we observed, future studies should explicitly assess the compositional differences in these two arthropod groups in communities separated by varying distances of forest and non-forest areas, with greater resolution at finer spatial scales, and across a more complete gradient of urban-ness.

The role of environmental filtering in shaping communities is demonstrated by the importance of habitat variation, in this case measured via proportion forest cover, for both arthropod groups. Corresponding to the importance of forest cover for arthropod community composition, the first principal component axis (PC1) is strongly associated with differences in forest cover for both foliage and ground arthropods. This finding aligns with the results of previous work, where differences in arthropod community composition are often associated with environmental differences between forested areas and urban areas [11,15,16,33]. Differences in community composition with forest cover, unrelated to diversity, emphasize how variation within the local environment shapes communities. In many cases, a smaller proportion of forest cover may be associated with smaller patch size, and smaller patches have greater edge density relative to interior habitat area. An increase in forest area could increase the suitable habitat for forest specialists, while decreasing the suitable habitat for edge specialists, thereby changing the composition of the arthropod community without necessarily impacting diversity. For foliage arthropods, two diet and habitat generalist families (Araneidae and Sclerosomatidae) are more abundant at plots with greater forest cover, while Coccinellidae, a family whose species largely specialize on aphids or scale insects and are often habitat specialists [34], is more abundant at plots with lower forest cover (Fig 4a). Among ground arthropods, wolf spiders (Lycosidae) and camel crickets (Rhaphidophoridae), both diet generalists but habitat specialists [35], are more abundant at sites with greater forest cover. (Fig. 4b). These differences in patterns of abundance based on the specialization of families aligns with previous studies suggesting that differences in community composition associated with the local environment tend to be driven by the habitat availability for both specialist and generalist taxa, where one group of specialists replaces another, or generalists replace specialists, depending on the types of habitat available in the local environment [36–38]. Taxa in urban environments tend to share traits that make them more suitable to those environments [39], and while our results do not provide an indication of which traits those may be, the response of both foliage and ground arthropod community composition to environmental traits associated with urbanization aligns with this pattern.

There were no predictors of differences in arthropod community composition that were significant for one metric of dissimilarity but not for the other, which is informative for researchers attempting to compare results between studies that used different metrics. Nevertheless, models with Euclidean distance as the response variable generally explained more variation in community composition than models with Jaccard dissimilarity as the response. This suggests that differences in the biomass of particular families, rather than the presence or absence of some families, were the primary drivers of differences in community composition resulting from both selection at the local level and dispersal limitation. The only exception to this pattern was the effect of resistance between plots on ground arthropod communities, where more variance was explained for Jaccard dissimilarity than Euclidean distance. This suggests that, after accounting for variation in the landscape, dispersal limitation primarily alters ground arthropod community composition by determining the presence or absence of particular families. Had we analyzed turnover at the level of species rather than families, it is possible that Jaccard dissimilarity might have had additional power to discriminate species-level habitat specialization that was otherwise unapparent.

Differences in the relevant environmental factors for foliage and ground arthropods further support the hypothesis that the relative abundance of generalists and different groups of specialists drive community composition. With forest cover most strongly predicting differences in ground arthropod communities, we see evidence that variation in communities depends on the abundances of taxa which specialize on particular habitats - in this case, primarily forested, urban, or edge environments. In the case of foliage arthropods, we see differences in community composition driven by the relative abundance of diet specialists, as well. While forest cover was predictive of foliage arthropod community variation, the dissimilarity of tree species on which arthropods were collected was a stronger predictor. Even within genera, different species of trees support distinct herbivorous arthropod communities [40], and these different herbivore communities may in turn shape arthropod predator communities, with greater tree diversity corresponding to greater spider abundance [41]. The importance of variation in the environment with regard to multiple factors - in this case, habitat and diet - reinforces the value of assessing drivers of community composition for groups which utilize different components of the same landscape. Furthermore, it demonstrates how modification of the environment can result in compounding changes to community composition when that modification occurs across multiple strata of an ecosystem.

Our work emphasizes the importance of both environmental filtering and dispersal limitation in shaping community composition, as well as the scale-dependence of these forces. For both groups included in this study, variables related to the local environment and dispersal between plots were strongly predictive of differences in the communities at those plots. Additionally, variation in the environment moderated both selection and dispersal: in the local environment, the proportion of different habitat types was important, while the variance in the landscape between sites, which altered the dispersal paths between plots, accounted for much more variation in arthropod communities than the straight-line distance between plots. These factors, taken together, highlight how local variation shapes the communities found across the landscape.

## Conclusion

Because urbanization can affect ecosystems in a variety of ways, it is essential that we investigate the roles of specific mechanisms in shaping communities. Here, we investigated how the local environment and the connectivity between plots structure arthropod communities in the foliage and on the ground. Environmental traits at each plot, particularly those associated with variation in habitat or food resources, strongly modified community composition, while the dispersal distance between plots was a much stronger predictor of differences in composition when it accounted for how landscape variation between plots modifies dispersal. Future work should focus on assessing whether these patterns occur across a broader range of spatial scales and at finer taxonomic resolutions, as well as further investigating how aspects of the landscape can moderate dispersal.

## Acknowledgements

We thank John Bruno and Steve Frank for their feedback on a previous draft and Jacob Fleenor for assistance with sample collection, and we acknowledge funding from NSF Macrosystems Biology grant EF-1702708.

## Supporting Information Captions

**Fig. S1: Abundance of arthropods by flight ability of species in each family.**

**Fig. S2: Scatterplots (a-c) of estimates of resistance of the shortest path between sampling plots using three different models of (d) how resistance values vary with land cover classes from the National Land Cover Database.** Low resistance implies a species can disperse easily across that land cover type while high values imply the land cover type is a barrier to dispersal.

